# Estimating yields of prenatal carrier screening and implications for design of expanded carrier screening panels

**DOI:** 10.1101/429050

**Authors:** Michael H. Guo, Anthony R. Gregg

**Affiliations:** Program in Medical and Population Genetics, Broad Institute of Harvard and MIT, Cambridge, MA 02142, USA; Department of Medicine, University of North Carolina Hospitals, Chapel Hill, NC 25779, USA; Department of Obstetrics and Gynecology, Baylor University Medical Center, Dallas, TX 75246, USA

**Keywords:** expanded carrier screening, exome sequencing, Mendelian disease, pathogenic variants, genetic counseling

## Abstract

**Purpose:** Prenatal genetic carrier screening can identify parents at risk of having a child affected by a recessive condition. However, the conditions/genes most appropriate for screening remain a matter of debate. Estimates of carrier rates across genes are needed to guide construction of carrier screening panels.

**Methods:** We leveraged an exome sequencing database (n=123,136) to estimate carrier rates across 6 major ancestries for 416 genes associated with severe recessive conditions.

**Results:** 36.5% (East Asian) to 65% (Ashkenazi Jewish) of individuals are variant carriers in at least one of the 416 genes. For couples, screening all 416 genes would identify 0.4-2.8% of couples as being at-risk for having a child affected by one of these conditions. Screening just the 47 genes with carrier rate > 1.0% would identify more than 85% of these at-risk couples. An ancestry-specific panel designed to capture genes with carrier rates > 1.0% would include 6 to 30 genes, while a comparable pan-ethnic panel would include 47 genes.

**Conclusion:** Our work guides the design of carrier screening panels and provides data to assist in counseling prospective parents. Our results highlight a high cumulative carrier rate across genes, underscoring the need for careful selection of genes for screening.

## Introduction

Prenatal genetic carrier screening (PGCS) has changed rapidly over the past few decades, driven by advances in technology, increased awareness of rare inherited conditions and their impact on families, and increased availability of treatments for inherited rare diseases. The paradigm has moved from screening only a limited number of variants (e.g., Δ508 in *CFTR*) in a small handful of conditions (e.g., *CFTR* for cystic fibrosis) in only high-risk populations (e.g., individuals of Caucasian ancestry) to screening many variants in a large number of genes across broad ancestry groups^1,2^. The latter screening paradigm has been called expanded carrier screening (ECS). ECS leverages next generation sequencing or high throughput genotyping to allow simultaneous and affordable assessment of genetic variation of a panel of genes. However, it is unclear which genes and variants should be included on ECS panels. Similarly, it is unclear whether individuals of all ancestries should receive the same screening panel or whether separate panels of genes should be tested for each ancestry.

The costs and potential benefit should be considered when deciding which genes should be included as part of an ECS panel. The cost of adding genes to a panel comes in two quantifiable forms. The first is the technical cost, which is becoming increasingly negligible in the era of next-generation sequencing ^3^. The second quantifiable cost comes from downstream interpretation and counseling. Adding genes to a panel will increase the number of variant carriers identified, which will necessitate additional counseling as well as confirmatory and follow-up testing, including testing of the patient’s partner. A third cost, which is very challenging to quantify, is the cost of anxiety to the patient. Identifying carriers of pathogenic variants from an ECS panel potentially increases stress and anxiety, though existing evidence suggests that there may not be much anxiety provoked by genetic testing ^4,5^. The benefit of adding genes to a panel is that it can increase identification of at-risk couples, which can inform pre-conception decisions and decisions regarding management of an established pregnancy.

Professional organizations including the American College of Obstetricians and Gynecologists (ACOG) and the American College of Medical Genetics and Genomics (ACMG) have made recommendations regarding the genes and conditions for which prenatal screening can or should be performed. ACOG recommended a lower limit carrier frequency (1/100) for screening conditions; however, this threshold is not based on empirical evidence ^6^. These recommendations have implications for financial stakeholders and public health given the approximately 4 million births in the United States annually. Therefore, additional data is needed to inform professional organization screening guidelines.

Cost and benefit of ECS cannot be determined until conditions appropriate for screening are better understood from a population perspective. In this study, we leveraged a population database with existing sequencing data to estimate the carrier rates for all severe recessive Mendelian conditions in order to inform the design and utility of ECS panels. We also sought to use this data to understand how the number of conditions screened influences the proportion of couples impacted.

### Materials and Methods

#### Study Population

We used data from gnomAD v2.0.2, which is comprised of summary-level data for 123,136 exome sequencing samples ^7^. The data was downloaded from http://gnomad.broadinstitute.org. For each variant in gnomAD, the allele count, allele number, and number of individuals who are homozygotes is provided for each of the ancestry groups. No individual-level data is available.

#### Variants and Genes Analyzed

To obtain a list of variants to analyze, we downloaded the ClinVar database ^8,9^. We used a version of the ClinVar database that was parsed in order to facilitate mapping of variants to genes and conditions^10^. We included all variants that are annotated as either “likely pathogenic” or “pathogenic” ^11^. A set of 40 variants (Table S1) were excluded from the analyses because they are common (MAF > 1.0% in at least one ancestry) and are of known low penetrance, or were excluded in a prior paper^12^.

We extracted a list of 924 genes previously annotated as being associated with severe Mendelian conditions ^13^. Among these 924 genes, we included the 416 genes that are annotated as acting in an autosomal recessive manner ^8^.

#### Estimation of gene carrier rate (GCR)

We first calculated the variant carrier rate (VCR) for each pathogenic or likely pathogenic variant.

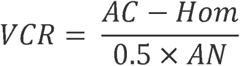

Here AC is the total allele count for the variant, Hom is the number of individuals who are homozygous for the variant, and AN is the total number of alleles analyzed for the variant.

The *GCR* for a gene *g* can then be estimated as:

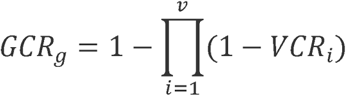

Here *VCR_i_* is the variant carrier rate for variant *i*, and *v* is the number of variants of interest in gene *g*.

These calculations were performed separately for each ancestry.

#### Estimation of cumulative carrier rate (CCR)

The *CCR* for a set of genes *s* can be estimated as:

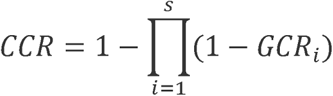

Here *GCR_i_* is the gene carrier rate for gene *i* in a set of *s* genes. These calculations were performed separately for each ancestry.

#### Estimation of at-risk couple rates

We also calculated the at-risk couple rate (ACR), which is the proportion of couples who each carry a likely pathogenic or pathogenic variant in the same gene. In contrast to prior work which calculated the proportion of affected offspring^12^, we eschewed this metric as our primary metric since the proportion of affected offspring is dependent on penetrance. The at-risk couple rate *ACR* can be estimated as:

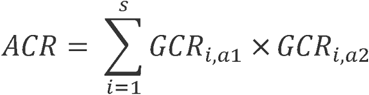

Here *GCR_i,a_*_1_ is the *GCR* for gene *i* in ancestry 1 and *GCR_i,a2_* is the *GCR* for gene *i* in ancestry 2. This calculation was performed for all *s* genes in a set of genes and across all unique pairwise combinations of *a*_1_ and *a*_2_.

#### Software

Processing of ClinVar data was performed using custom Python scripts (v2.7) and Bcftools v1.2^14^ Analyses and plotting was performed using R v3.1.

## Results

### Study Overview

In this study, we first characterized the variant carrier rates (VCR), which is the proportion of individuals who carry a given variant, across a set of 416 genes previously ascribed as being associated with a severe autosomal recessive disorders ^13^. We calculated these VCRs using data from an exome sequencing database (gnomAD) comprised of 123,136 samples^7^. Importantly, although not truly a population-based cohort, the individuals included in gnomAD have no known history of a severe Mendelian condition. These individuals were sequenced across many different sequencing platforms and at different centers, but the sequencing data underwent uniform quality control and joint variant calling.

The samples in gnomAD are distributed across 7 major ancestries: African/African American (AFR, n=7652), Hispanic (AMR, n=16791), Ashkenazi Jewish (ASJ, n=4925), East Asian (EAS, n=8624), Finnish (FIN, n=11150), non-Finnish European (NFE, n=55860), and South Asian (SAS, n=15391). We did not include the Finnish in this study since they comprise a small proportion of the US population. We also calculated variant frequencies for a composite US sample, which takes the variant frequencies from gnomAD and scales them by 2016 US census data^15^.

We then used these VCRs to generate estimates of gene carrier rates (GCRs) for each gene, which is the estimated proportion of individuals who carry one or more pathogenic or likely pathogenic variants in that gene. Using these GCRs, we also calculated cumulative carrier rates (CCRs) for various sets of genes. These sets of genes are based on thresholds (e.g., genes having a GCR greater than a given percentage) and are meant to simulate hypothetical carrier screening panels. Please see Figure 1 for a depiction of our workflow.

**Figure 1:**
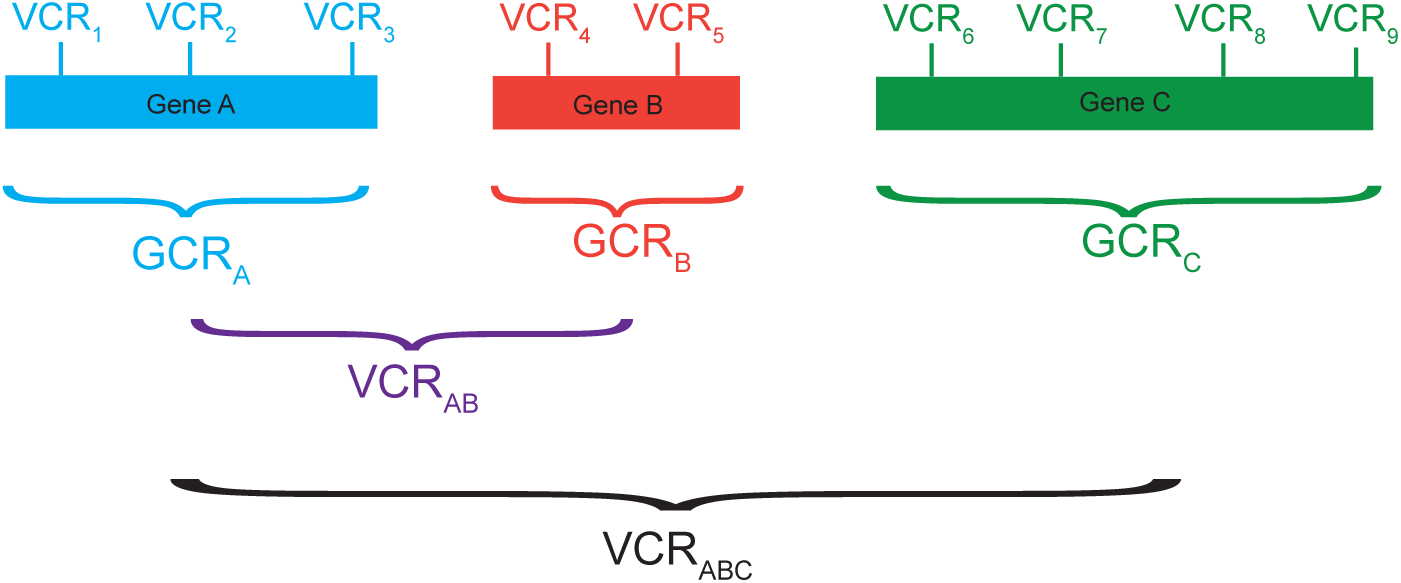
Scheme for paper. On top, are shown three genes: A (blue), B (red), and C (green). Each gene has between 2-4 variants, each with an associated variant carrier rate (VCR). The gene carrier rate (GCR) for each gene is then the probability that an individual will carry at least one variant in the given gene. Finally, cumulative carrier rates (CCRs) are shown for two sample panels of genes. CCR_AB_ (purple) is the CCR for a hypothetical panel of genes containing genes A and B. CCR_ABC_ (black) is the CCR for a hypothetical panel containing genes A, B, and C.

### Carrier rates by gene

The GCR for each of the 416 genes associated with severe recessive Mendelian conditions for each of the 6 ancestries as well as the US composite are listed in Table S2. For illustrative purposes, Figure 2 shows the ten highest GCRs for each of the 6 ancestry groups. Across ancestries, the highest GCR for a single gene is 12.0% for *HBB* (encoding the hemoglobin beta chain, mutations in which cause hemoglobinopathies including sickle cell anemia) in AFR. In each ancestry, the carrier rates rapidly decline. For example, in ASJ, only 30 of the 416 genes had a carrier rate > 1%. In contrast, in AMR, only 6 of the 416 genes had a GCR > 1%.

**Figure 2:**
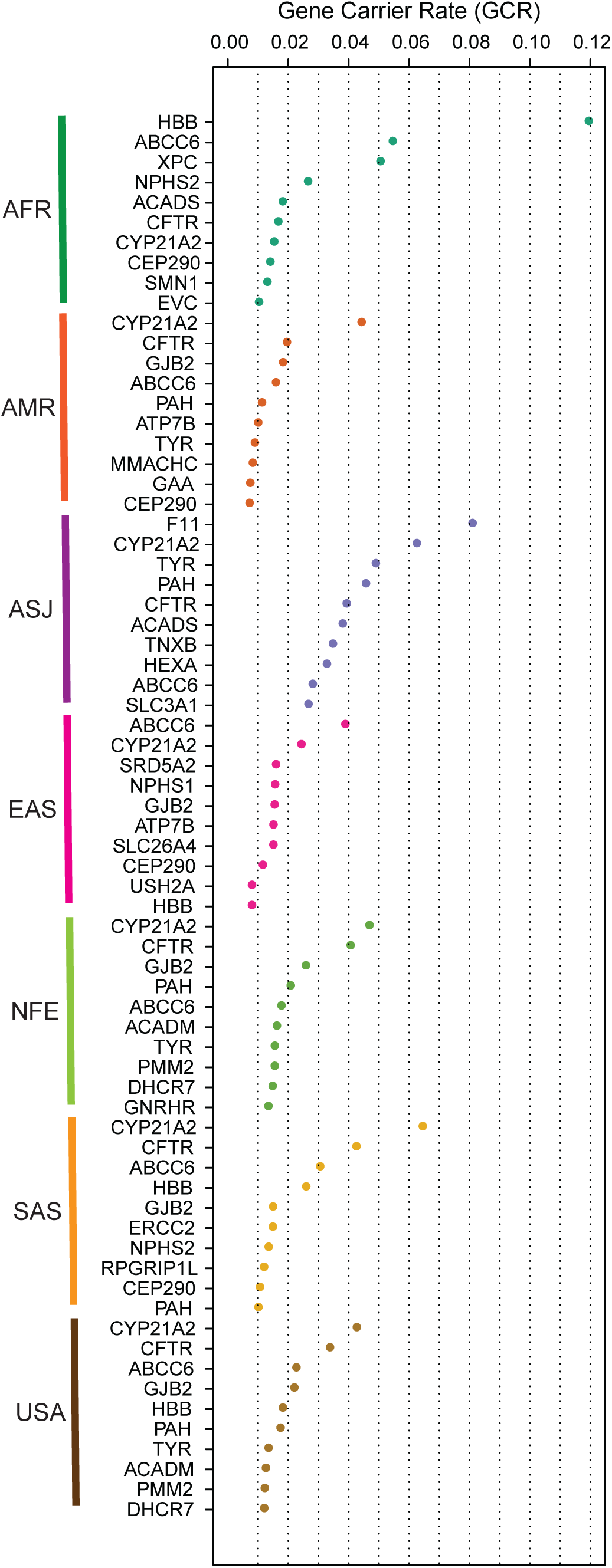
Gene carrier rates (GCRs) for the top ten genes for each ancestry. The genes are listed on the vertical axis, and the GCRs are shown on the horizontal axis.

In Figure S1, we show the top 30 genes in terms of GCR in any ancestry and show the carrier rates across ancestries for each of these genes. As can be seen, some genes such as *HBB* or *F11* are relatively restricted to a single population (AFR and ASJ respectively). In contrast, other genes such as *CYP21A2* have high GCRs across many ancestries.

### Cumulative Carrier Rates

We ranked each gene in descending order by its GCR in the respective ancestry and plotted the CCRs as increasing numbers of genes are screened. As can be appreciated in Figure 3A, there is initially a rapid increase in the CCR, reflecting a small number of genes with high GCRs. This is then followed by a long tail of genes which contribute asymptotically to the CCR. For example, in ASJ, 90% of the CCR is contributed by the 45 top ranking genes with the highest GCR (out of 416 total genes).

**Figure 3:**
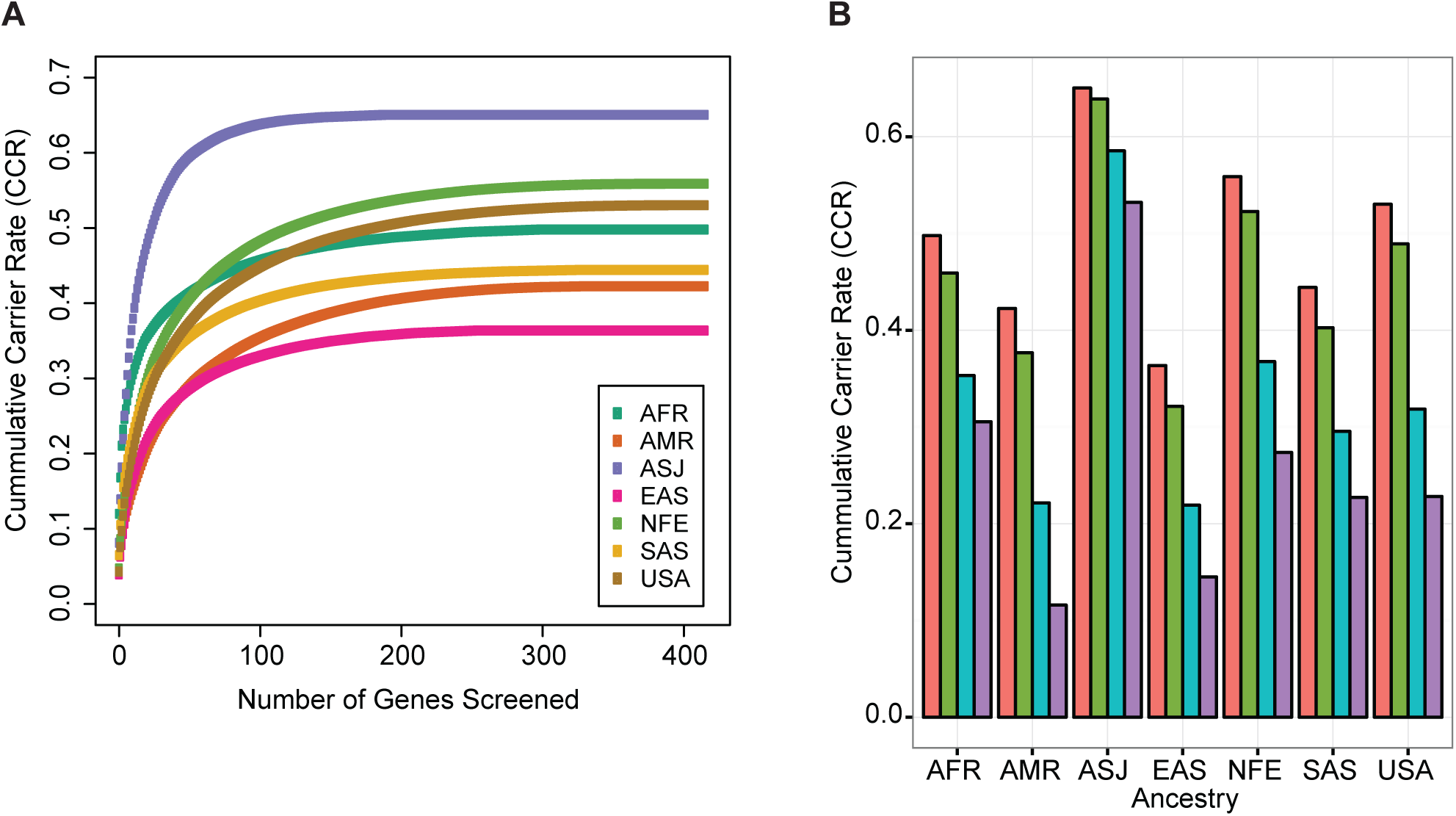
A) Cumulative carrier rates (CCRs) for each ancestry as additional genes are screened. Genes are ranked for each ancestry in descending order based on GCR for that ancestry. B) CCR when screening all 416 genes (pink), only genes with GCR > 0.1% (green), > 0.5% (blue), or > 1.0% (purple).

We next calculated CCRs for various sets of genes, which are delineated to reflect the hypothetical construction of an ECS panel. We first examined the CCR of all 416 severe recessive genes. As can be appreciated in Figure 3B, there were very high CCRs for this set of 416 genes in all ancestries (red bars). At the extreme, in ASJ, the CCR across all 416 genes was 65.0%. The lowest CCR for the 416 genes was in EAS at 36.5%.

The American College of Obstetrics and Gynecology (ACOG) recently suggested that genes with GCR > 1.0% are appropriate for ECS. We thus examined how screening only genes with GCR > 1% in the respective population would affect the CCR. We found that setting this >1% threshold drastically reduces the CCR by 20.1-55.8% (purple bars) (Figure 3B). We also show CCRs if only genes with GCR > 0.5% (blue bars) or > 0.1% (green bars) are included (Figure 3B). We note that these results simulate the yields that would be derived from using a hypothetical ancestry-specific ECS panel, where only genes with GCR greater than some threshold based on that ancestry are included on the panel.

### Design of Pan-ethnic Panels

We next examined the number of genes that meet the 1.0% GCR threshold in the respective ancestry. We found that in ASJ, 30 genes had GCR > 1.0 %, while in AMR, only 6 genes had GCR > 1.0% (Figure 4A, blue bars). Lowering the GCR threshold to > 0.5% (green bars) or > 0.1% (pink bars) greatly increases the number of genes that need to be screened.

**Figure 4:**
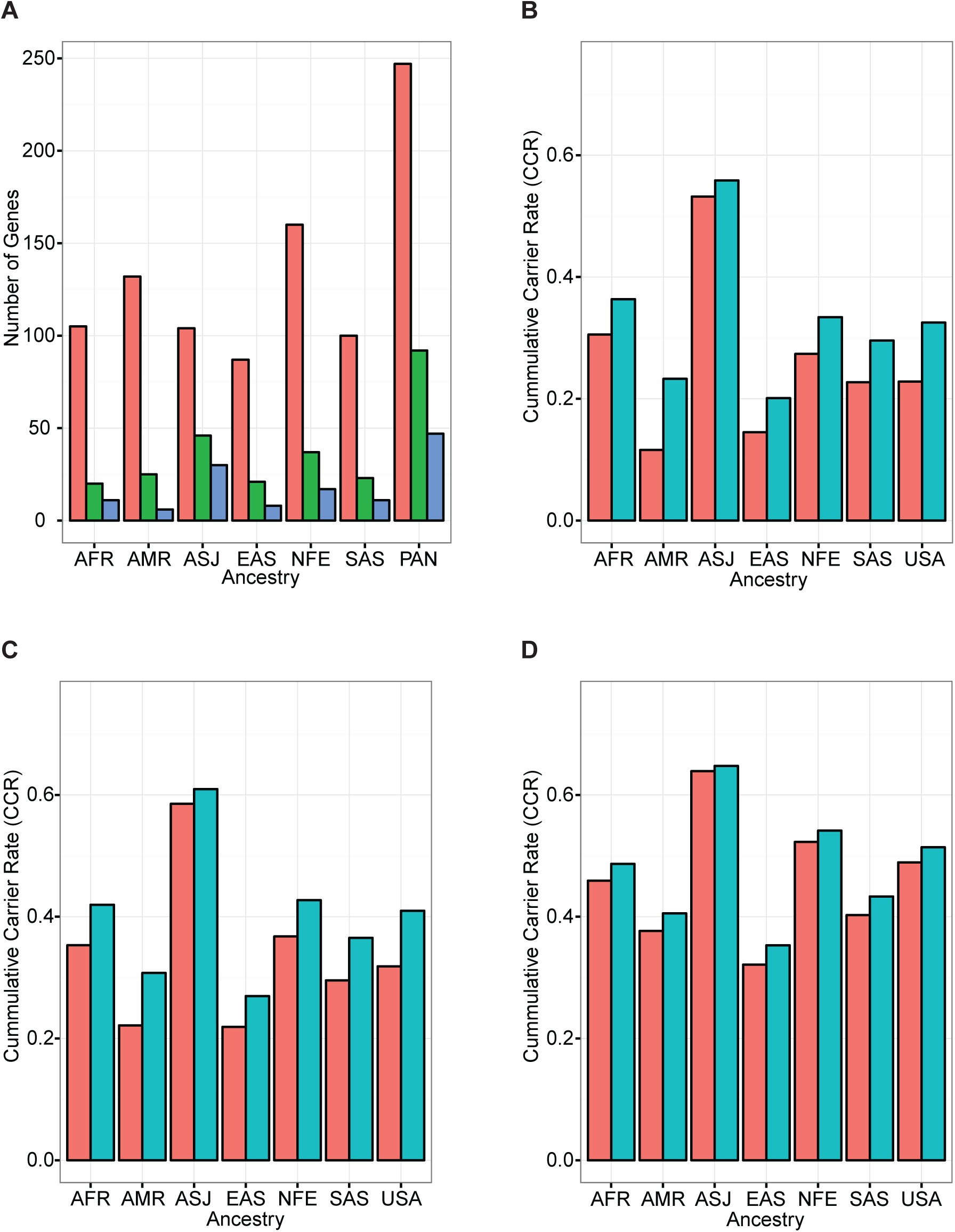
A) Number of genes with GCR > 0.1% (pink), > 0.5% (green), or > 1.0% (purple) for each of six ancestries or for a hypothetical pan-ethnic panel (PAN). B-D) CCR for each ancestry using an ancestry-specific or pan-ethnic panel comprised of genes with GCR > 1.0% (B), > 0.5% (C), or > 0.1% (D).

The previous analyses (Figure 3) defined hypothetical ECS panels using GCR thresholds in each respective ancestry and are thus ancestry-specific. However, it may be desirable to apply a single panel across all ancestries (i.e., pan-ethnic panel). We thus sought to evaluate the performance of a hypothetical pan-ethnic panel. We calculated the number of genes that would need to comprise a pan-ethnic panel such that all genes with a GCR greater than a threshold in *any one* of the individual ancestries would be included. We found that pan-ethnic panels greatly increase the number of genes needed to be screened. For example, in order to include all genes with GCR > 0.1% in any component ancestry, 247 genes would need to be on the pan-ethnic panel. In contrast, there was a range of 87 (EAS) to 160 (NFE) genes with GCRs > 0.1% in each of the individual ancestries.

We next calculated the CCR for each pan-ethnic panel and compared it to ancestry-specific panels (Figure 4B-D). Figure 4B shows the CCRs for a hypothetical pan-ethnic panel (blue bars) comprised of genes with GCR > 1.0% as compared to similar ancestry-specific panels (red bars). Parallel analyses are shown for panels designed to capture genes with GCR > 0.5% (Figure 4C) and for GCR > 0.1% (Figure 4D).

### At-risk couple rate

We examined the at-risk couple rate (ACR) for all 416 genes, which is the probability that both the mother and father are carriers for pathogenic or likely pathogenic variants in the same gene. Unsurprisingly, the highest ACRs for intra-ancestry couples was in ASJ (280 out of 10,000 couples) and the lowest was for AMR (47 out of 10,000 couples (Figure 5A). The inter-ancestry rates ranged from 40 out of 10,000 EAS/NFE couples, to 105 out of 10,000 NFE/ASJ couples.

**Figure 5:**
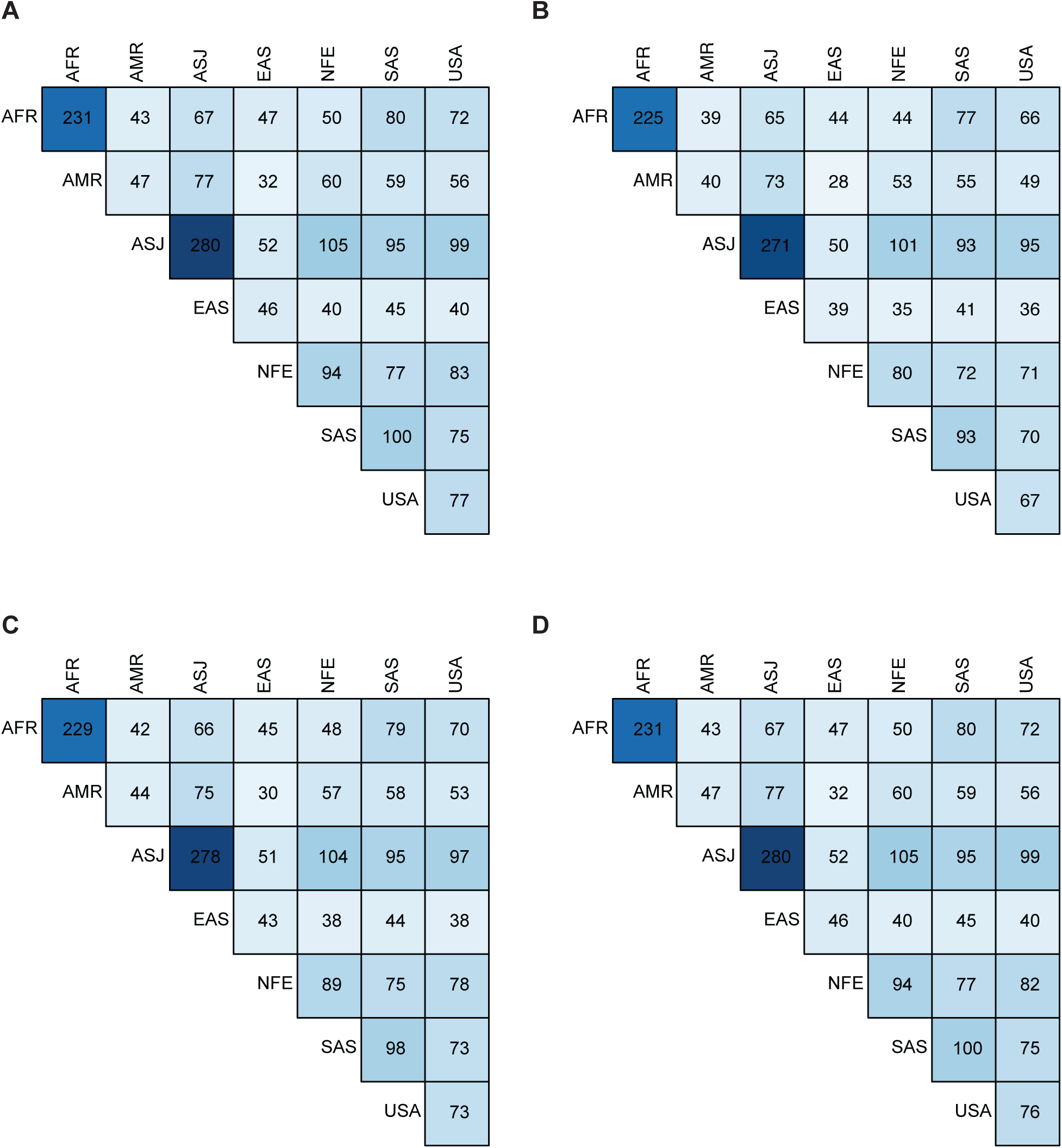
A) Number of couples (out of 10,000) that are at-risk when screening all 416 genes associated with a severe recessive condition. The ancestries are indicated on the top and the diagonal. The boxes in the diagonal shows rates for couples where both individuals are of the same ancestry, and other boxes are for couples where individuals are of different ancestries. B-D show similar data, except when considering a hypothetical pan-ethnic panel comprised of genes with GCR > 1.0% (B), > 0.5% (C), or > 0.1% (D).

We also calculated the ACR when considering only genes with GCR > 1.0% in any population (Figure 5A), GCR > 0.5% (Figure 5C), or GCR > 0.1% (Figure 5D), reflecting the ACR that would result from the application of various hypothetical pan-ethnic panels highlighted in Figure 4. Strikingly, the yields in terms of ACR for screening just the 47 genes with GCR > 1.0% are very close to the yields of screening all 416 genes (compare Figure 5A with Figure 5B). In fact, screening just these 47 genes would identify at least 85% of the at-risk couples that screening all 416 genes would identify.

## Discussion

In this study, we have leveraged a large-scale exome sequencing database to estimate the GCR for each gene associated with a severe recessive condition. We also estimated the CCRs for various sets of genes to simulate the yields of hypothetical ECS panels at various specified carrier frequency thresholds. We found very high CCRs across ancestries, including 65% in ASJ. We found that the proportion of at-risk couples was comparatively lower, ranging from 0.4-2.8% of couples.

Our study has important implications for the design of ECS panels. First, we observed that CCRs increase asymptotically as additional genes are screened. This suggests that a small number of genes accounts for the majority of carrier rates across genes, which is followed by a long tail of genes with a relatively low yield. Second, the CCRs that we observed were very high, as high as 65% in ASJ. This suggests that a large fraction of individuals who undergo PGCS will be a carrier for a pathogenic or likely pathogenic variant in a severe recessive Mendelian condition. This introduces the possibility of substantial cost from downstream follow-up when an individual screens positive. This underscores the need for careful consideration of which genes should be screened. Third, our study provides insight into the efficacy of pan-ethnic panels. At more stringent carrier rate thresholds such as GCR > 1.0 %, constructing a pan-ethnic panel would require relatively few additional genes as compared to an ancestry-specific panel. In contrast, at less stringent thresholds such as GCR > 0.1%, a pan-ethnic panel would require a much larger number of genes than any ancestry-specific panel. Despite much greater numbers of genes needed to be included on a pan-ethnic panel as compared to an ancestry-specific panel, the incremental yields in terms of CCR were minimal.

Our paper highlights a critical insight—although the GCRs are very high for any panel of genes, the proportion of at-risk couples are comparatively much lower. For example, if all 416 genes associated with severe recessive conditions are screened in ASJ, which has the highest CCR (65%) among ancestries analyzed, 280 out of 10,000 couples (2.8%) would carry a likely pathogenic or pathogenic variant in the same severe recessive disease gene. For other ancestries, the proportion of at-risk couples is as low as 40 out of 10,000 couples (0.4%). The number of affected offspring is even lower than that: assuming full penetrance, 10 to 70 per 10,000 fetuses would be affected. The ACR for couples of different ancestries (inter-ancestry) is lower than the ACR for couples of the same ancestry (intra-ancestry). In fact, in nearly all cases, the ACR for an inter-ancestry couple is lower than the ACR for either component ancestry. As an example, the ACR across all 416 genes for ASJ/ASJ couples is 2.8%, and the ACR for AFR/AFR couples is 2.3%. However, the ACR for an ASJ/AFR couple is only 0.67%. This is a reflection of how the GCRs for many genes are relatively specific to a single population (Figure S1). For example, for AFR, the top gene is *HBB* (associated with various hemoglobinopathies; GCR=12.0%), but *HBB* has a GCR of just 0.020% in ASJ. In contrast, the top gene for ASJ is *F11* (associated with Factor XI deficiency; GCR=8.1%), but *F11* has a GCR of just 0.010% in AFR. For these genes where the GCR is relatively ancestry-specific, the rate of both individuals of a couple with discordant ancestries each carrying a variant will be very low.

Strikingly, we found that screening just a small handful of genes captures the vast majority of the at-risk couples from screening all 416 genes. For example, for an ASJ/ASJ couple, 2.80% of couples would be identified as being as at-risk when screening all 416 genes. However, screening just the 47 genes with GCR>1.0% in any ancestry would identify 2.71% of couples as being at-risk. Screening the 92 genes with GCR > 0.5% would identify 2.78% of couples, and screening the 247 genes with GCR>0.1% would identify nearly all 2.8% of at-risk couples. This suggests that lowering the threshold for genes that should be screened greatly increases the number of genes that would need to be screened, but results in a modest increase in the CCR and a miniscule increase in the ACR.

This study is the first to comprehensively evaluate carrier rates for recessive conditions. Haque et al. performed similar analyses for a commercial ECS panel. However, their study was limited in that it only examined 94 genes^12^. Also, many of the samples were analyzed using targeted genotyping panels, and thus was not comprehensive in terms of the variants examined in the targeted genes. Other studies have been more comprehensive in terms of variants and genes analyzed, but have been limited to only a few hundred of individuals analyzed^16–18^. Thus, our study provides a comprehensive and large-scale examination of carrier rates for many genes associated with recessive disorders.

There are several important limitations to our study. First, we only examined coding variants in autosomal recessive genes. Our study cannot be extended to genes acting through alternative modes of inheritance, such as X-linked recessive. We also could not reliably examine more complex forms of genetic variation, such as CNVs (e.g., deletions in *SMN1* causing spinal muscular atrophy) ^19^. Therefore, our data underestimates carrier rates for some genes. Second, characterizing the severity of disorders associated with genes in determining appropriateness for screening is a challenging task. Here, we used a previously published list of 416 genes associated with recessive conditions that were ascribed as being severe ^13^. However, what comprises a condition worthy of screening in terms of phenotypic features is a challenging technical and ethical question that is beyond the scope of this work. As a related issue, many of these genes might have lower penetrance and/or variable expressivity and therefore might not be appropriate for screening. Third, we chose to only include variants annotated as likely pathogenic or pathogenic in ClinVar ^8,9,11^. Some of these variants are actually benign, while there may be others that we did not include that are actually pathogenic ^20,21^. This is an issue that is well beyond the scope of this work.

In summary, our work informs the selection of genes that should be considered for screening in PGCS and provides an important foundation for policy makers as they consider the implications of pan-ethnic application of PGCS panels. Our data also facilitates counseling of patients as they consider PGCS.

## Acknowledgements

We thank the individuals who contributed their samples to the gnomAD database, as well as the gnomAD team for compiling the resource and sharing it.

## Supplementary Figure Legends

Figure S1: Gene carrier rates (GCRs) for the top 30 genes based on highest GCR in any ancestry. The x-axis shows the genes, ranked from highest to lowest GCR. The y-axis shows the GCR. Each gene has six points, representing the GCR for each of six ancestries.

## Supplementary Table Legends

Table S1: Variants that were excluded from analyses. Coordinates are based on human reference genome GRCh37 and annotations were taken from ClinVar.

Table S2: Gene carrier rates (GCR) for each of 416 genes associated with severe, recessive conditions. Columns 2-7 show the GCR for each of six ancestries. Column 8 shows the GCR for the US composite. Column 9 (MAX_GCR) is the maximum GCR among the 6 individual ancestries. Column 10 shows the conditions associated with that gene. Genes highlighted in green have a GCR> 1.0% in at least one ancestry. Genes highlighted in yellow have GCR between 0.5% and 1.0%. Genes highlighted in grey have GCR between 0.1% and 0.5%. Thus, a pan-ethnic panel with threshold of GCR>1.0% would include all genes highlighted in green; a pan-ethnic panel with threshold of GCR>0.5% would include all genes highlighted in green or yellow; a pan-ethnic panel with threshold of GCR>0.1% would include all genes highlighted in green, yellow, or grey.

